# Therapeutic administration of mouse mast cell protease 6 improves functional recovery after traumatic spinal cord injury in mice by promoting remyelination and reducing glial scar formation

**DOI:** 10.1101/2022.12.08.519463

**Authors:** Tim Vangansewinkel, Stefanie Lemmens, Assia Tiane, Nathalie Geurts, Dearbhaile Dooley, Tim Vanmierlo, Gunnar Pejler, Sven Hendrix

**Author notes:** **Corresponding author:** Sven Hendrix, MD, PhD, Medical School Hamburg, Am Kaiserkai 1, 20457, Hamburg, Germany.

## Abstract

Traumatic spinal cord injury (SCI) most often leads to permanent paralysis due to the inability of axons to regenerate in the adult mammalian central nervous system (CNS). In the past, we have shown that mast cells (MCs) improve the functional outcome after SCI by suppressing scar tissue formation at the lesion site via mouse mast cell protease 6 (mMCP6). In this study, we investigated whether recombinant mMCP6 can be used therapeutically to improve the functional outcome after SCI. Therefore, we applied mMCP6 locally via an intrathecal catheter in the subacute phase after a spinal cord hemisection injury in mice. Our findings showed that hind limb motor function was significantly improved in mice that received recombinant mMCP6 compared to the vehicle-treated group. In contrast to our previous findings in mMCP6 knockout mice, the lesion size and expression levels of the scar components fibronectin, laminin, and axon-growth-inhibitory chondroitin sulfate proteoglycans were not affected by the treatment with recombinant mMCP6. Surprisingly, no difference in infiltration of CD4+ T cells and reactivity of Iba-1+ microglia/macrophages at the lesion site was observed between the mMCP6-treated mice and control mice. Additionally, local protein levels of the pro- and anti-inflammatory mediators IL-1β, IL-2, IL-4, IL-6, IL-10, TNF-α, IFNγ, and MCP-1 were comparable between the two treatment groups, indicating that locally applied mMCP6 did not affect inflammatory processes after injury. However, the increase in locomotor performance in mMCP6-treated mice was accompanied by reduced demyelination and astrogliosis in the perilesional area after SCI. Consistently, we found that TNF-a/IL-1B-astrocyte activation was decreased, and that oligodendrocyte precursor cell (OPC) differentiation was increased after recombinant mMCP6 treatment *in vitro*. Mechanistically, this suggests effects of mMCP6 on reducing astrogliosis and improving (re)myelination in the spinal cord after injury. In conclusion, these data show for the first time that recombinant mMCP6 is therapeutically active in enhancing recovery after SCI.

## Introduction

Spinal cord injury (SCI) is an incurable chronic condition that typically results in functional impairment in locomotion, and loss of sensation below the lesion site. It can also cause neuropathic pain, spasticity, and incontinence (1-3). The pathology of traumatic SCI is characterized by two distinct phases: first, a primary traumatic insult that rapidly destroys neuronal cells and axons (4, 5), followed by a wave of secondary injury processes that cause further tissue damage and exacerbate the neurological deficits (6, 7). These processes include the initiation and propagation of a robust pro-inflammatory response and the formation of a glial and fibrous scar at the lesion (8-10). Several biochemical and cellular changes are triggered after injury that affect the local environment in the spinal cord. These changes generally start with the activation and influx of inflammatory cells such as microglia, macrophages, and T cells into the damaged tissue, and the local secretion of inflammatory mediators (e.g., TNF-α, IL-1β, MCP-1 and matrix metalloproteinases (MMPs)) (11, 12). As an essential part of the glial scar, reactive astrocytes can exert harmful effects on the functional outcome via several mechanisms: [1] production of scar matrix components that are inhibitory to axon regeneration (e.g. chondroitin sulfate proteoglycans (CSPGs), tenascins, and ephrins), [2] formation of a glia limitans that acts as a physical barrier, and [3] by releasing pro-inflammatory and toxic mediators. The latter can induce secondary tissue damage (8, 13-16), although beneficial functions have also been reported. Neurons and oligodendrocytes are especially vulnerable to these toxic insults, leading to further injury of axons that were left intact by the initial trauma and demyelination (17, 18). In addition, mediators (e.g., MicroRNA-26b) that stimulate the differentiation of oligodendrocyte precursors cells (OPCs) towards myelinating oligodendrocytes are downregulated (19). However, these processes are necessary to contribute to remyelination and functional recovery and are susceptible to therapeutic intervention (20-22).

Mast cells (MCs) are essential effector cells of the immune system, and play a major role in neurotrauma (23). Their most distinguishing morphological feature is the high content of electron-dense lysosome-like secretory granules that occupy a major part of the cytoplasm in mature MCs. Several preformed and preactivated mediators are stored in these granules, such as IL-4, nerve growth factor (NGF), and various MC-specific proteases (tryptases, chymases, and carboxypeptidase A) (24-27). When MCs are triggered to degranulate, these compounds are released into the extracellular environment and can have marked physiological and pathophysiological effects. Moreover, we have shown that MCs protect the CNS after traumatic injury via different mechanisms: by suppressing harmful inflammatory reactions via mouse mast cell protease 4 (mMCP4) that cleaves pro-inflammatory mediators (28, 29) and by reducing scar tissue formation via mMCP6 (30). MMCP6 is a serine protease with trypsin-like cleavage specificity and it is the most abundant protease stored and secreted by ‘connective tissue-type’ MCs (mMCP6 corresponds to β-tryptase in human MCs) (24, 31, 32).

In the current study, we explored the therapeutic potential of mMCP6 after traumatic SCI. We show that recombinant mMCP6 improves hind limb motor recovery when it is repeatedly administered in the subacute phase after a hemisection SCI in mice. This improvement in functional outcome was also associated with a reduction in demyelination and astrogliosis at the lesion site. Moreover, recombinant mMCP6 suppressed inflammation-induced astrogliosis *in vitro*, and it also stimulated the differentiation of oligodendrocyte precursor cells *in vitro*. In conclusion, these data show for the first time that mMCP6 can be used as a potential therapy to restore lost functions after SCI by diminishing detrimental glial scarring and boosting remyelination.

## Materials and methods

### Animals

All *in vivo* experiments were performed using 10-to 12-week-old female wild-type (WT) C57BL/6j mice (Janvier). Animals were housed in a conventional animal facility at Hasselt University under regular conditions, i.e. in a temperature-controlled room (20 ± 3°C) on a 12 hour light-dark schedule and with food and water *ad libitum*. Experiments were approved by the local ethical committee of Hasselt University and were performed according to the guidelines described in Directive 2010/63/EU on the protection of animals used for scientific purposes.

### Intrathecal catheter implantation

An intrathecal catheter was implanted as previously described (33, 34). Briefly, mice were anesthetized by intraperitoneal injection of ketamine (100 mg/kg, Ketalar®, Pfizer) and xylazine (12 mg/kg, Rompun®, Bayer) and a small incision was made in the skin at the base of the skull. Using curved forceps, muscles (m. interscutularis) connected to the external occipital crest were detached and retracted to expose the atlanto-occipital membrane, and also the underlying medulla oblongata becomes visible at this point. The head of the mouse was flexed and an opening was made in the membrane with a small needle (32 gauge), leading to the leakage of cerebrospinal fluid (CSF) from the cisterna magna. A mouse intrathecal catheter (Alzet^®^, DURECT Corp.) was carefully inserted via this opening into the intrathecal space at the midline, dorsal to the spinal cord. It was passed caudally (∼2.5 cm) into the subarachnoid space with the distal part of the catheter tip lying in close proximity to the lesion area (see illustration **Fig. 1A**). Tissue glue (Histoacryl^®^, Braun) was used to fix the catheter to the surrounding bone and muscle, and the catheter was externalized on the proximal side via a separate opening in the skin. Afterwards, the skin was sutured and a local subcutaneous injection of buprenorphine (0.1 mg/kg, Temgesic®, Schering Plough) was given as a post-operative pain treatment. To improve recovery after the surgery, an antisedative (0.5 mg/kg, Antisedan®, Orion Pharma) was administered intraperitoneally and mice were placed in a temperature-controlled chamber (33 °C) until thermoregulation was established. Afterwards, mice were housed individually and were allowed to recover for 7 days before SCI was induced. Mice that displayed neurological abnormalities and paralysis after the catheter implantation were excluded from the study (∼30% of the population).

**Figure 1.**
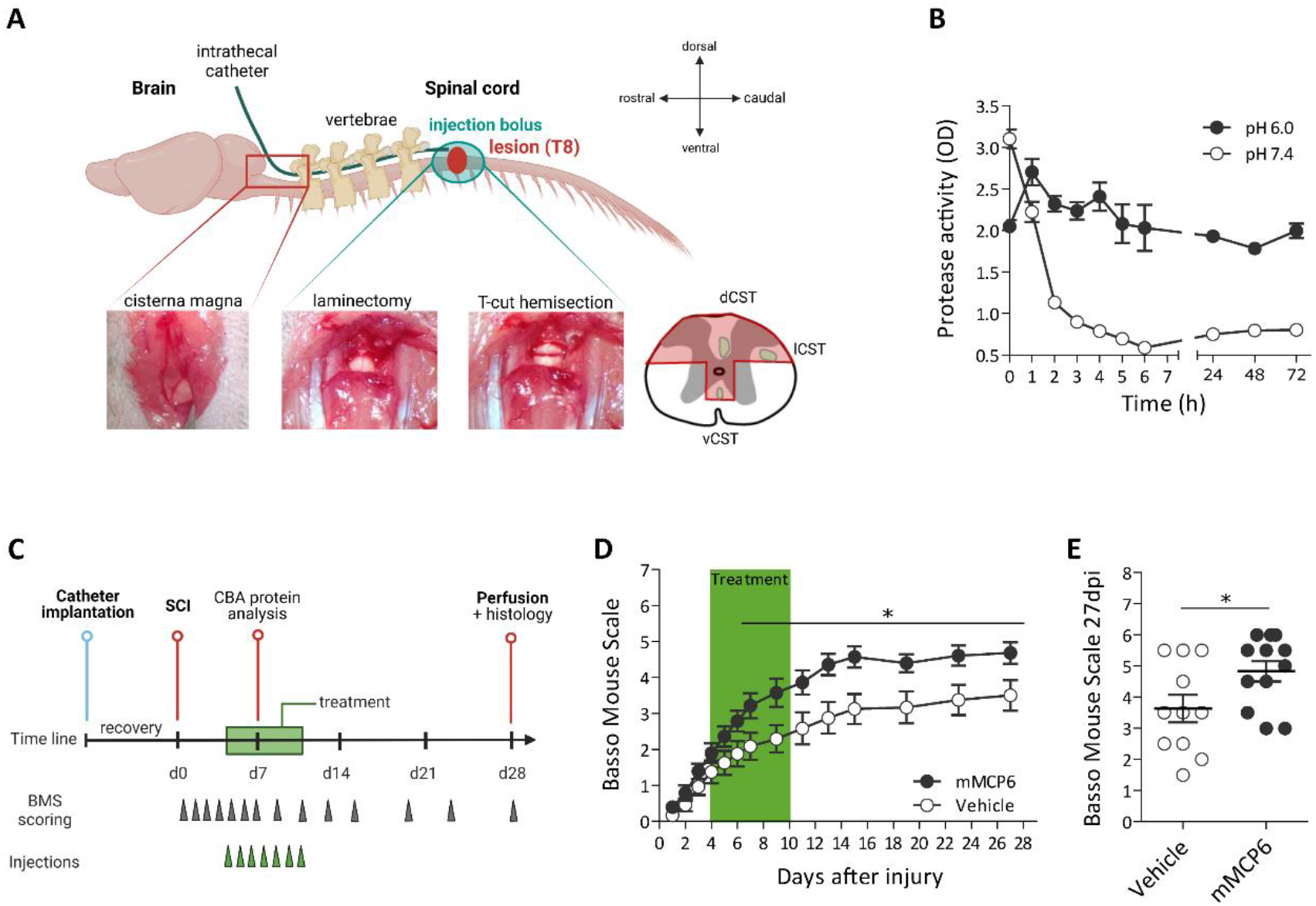
Local intrathecal delivery of recombinant mMCP6 in the subacute phase after SCI enhanced hind limb locomotor function in mice. **(A)** Illustration of the intrathecal cathether implantation via the cisterna magna and induction of a bilateral “T-cut” hemisection injury. Created with BioRender.com. More details are provided in the Material and Methods section. vCST: ventral corticospinal tract, dCST: dorsal corticospinal tract, lCST: lateral corticospinal tract. **(B)** The proteolytic stability of recombinant mMCP6 was determined at pH 7.4 and pH 6.0 (optimum for mMCP6 activity). Under physiologial conditions, the catalytic activity of recombinant mMCP6 decreased rapidly. **(C)** Time line of the experimental setup and protocol. SCI was induced in WT C57BL/6j mice and they received either recombinant mMCP6 or an inactive vehicle control, daily from 4 dpi until 7 dpi. Recovery of hind limb motor function was assessed using the Basso Mouse Scale (BMS). **(D)** Mice that received mMCP6 performed significantly better (improved hind limb function) compared to the vehicle control group, starting from 7 dpi until 27 dpi. Data are presented as mean values ± SEM. mMCP6: n = 12; vehicle: n = 11. **(C)** Individual BMS scores at 27 dpi. Two independent experiments performed under the same conditions. * p < 0.05

### Spinal cord hemisection injury

A “T-cut” hemisection injury was performed as previously described (29, 35). Briefly, mice were anesthetized and a partial laminectomy was performed at thoracic level 8 to expose the spinal cord. Next, a bilateral hemisection injury of the spinal cord was induced by using iridectomy scissors to transect left and right dorsal funiculus, the dorsal horns and additionally the ventral funiculus. This lesion typically results in a complete transection of the dorsomedial and ventral corticospinal tract, but it also impairs several other descending and ascending tracts (e.g. raphespinal tract) (**Fig. 1A**). Afterwards, the overlying muscles were sutured and the back skin was closed with wound clips (Autoclip®, Clay-Adams Co., Inc.). The following drugs were given after the operation: glucose solution (20%) to partially compensate for any blood loss during surgery, buprenorphine (0.1 mg/kg, Temgesic®) as a post-operative pain treatment and an antisedative (0.5 mg/kg, Antisedan®) to improve recovery after the surgery. All mice were placed in a temperature-controlled chamber (33°C) until thermoregulation was established and bladders were emptied manually until a spontaneous return of the micturition reflex. To standardize our model, all operations were performed by a trained experimenter and all mice were operated in the same way; animals that displayed abnormalities during the surgery were excluded from the study (e.g. substantial variations in morphology of the spinal blood vessels, unusual strong bleeding and abnormal response to the anesthesia).

### Production recombinant mMCP6 and drug administration

Recombinant mMCP6 was produced in an eukaryote expression system by using 293 EBNA Human Endothelial Kidney cells as described previously (30, 36). The proteolytic stability of recombinant mMCP6 was determined and verified under physiological conditions (i.e. 37°C, pH 7.4) by using the chromogenic substrate S-2288 (Chromogenix) (30). mMCP6 was inactivated by heating it at 100 °C for 10 min to generate an inactive vehicle, which was used as a control condition in this *in vivo* study. During drug administration, mice were immobilized with a volatile anesthetic (2% isoflurane; carrier gas O_2_). They were injected daily via the implanted catheter from 4 days post injury (dpi) until 10 dpi (subacute phase after injury), using a Hamilton syringe (#1701, Hamilton Co.) with either recombinant mMCP6 or with a proteolytically inactive vehicle (5 µl / 500 ng) that was used as a control. An additional injection of 5 µl PBS (pH 6.0) was carried out to ensure a proper and complete intrathecal diffusion of the drug. Treatments were coded, and evaluators were blinded to the treatment group.

### Locomotion testing

Locomotor recovery of the animals was assessed using the Basso Mouse Scale (BMS) (37, 38). The BMS is a 10-point locomotor rating scale (9 = normal locomotion; 0 = complete hind limb paralysis), in which mice are scored by an investigator blinded to the experimental groups, and which is based on hind limb movements made in an open field during a 4 min interval. Mice were scored daily during the first week, then they were scored every second day in the second week after injury and from the start of the third week until the end of the observation period (28 dpi), mice were examined every fourth day. We applied the following exclusion criteria: mice that still had a BMS score of zero at the end of the observation period (28 dpi) or mice that had a significantly higher score immediately after the operations were excluded from the study. In addition, mice that display abnormalities (e.g. pain behavior, infections) or dramatically lose weight (> 10% of body weight) during the experiment would have been excluded from the study. However, none of the mice showed these symptoms in our study.

### Immunohistochemistry and quantitative image analysis

At 28 dpi, mice received an overdose of pentobarbital (100 mg/kg, Nembutal®, Ceva) and were transcardially perfused with Ringer solution (1.98 mM NaCl, 44 mM KCl, 27 mM CaCl_2_, 5.8 mM NaHCO_3_, 70 mM NaNO_2_ and 280 mM phenol red in MilliQ) containing heparin (LEO Pharma), followed by 4% paraformaldehyde in phosphate-buffered saline (PBS, pH 7.4). Next, 14 µm thick saggital tissue sections were cut and immunohistochemical stainings were performed. Spinal cord sections were blocked with 10% normal goat serum and permeabilized with 0.05% Triton X-100 in PBS for 30 min at room temperature (RT). Then, the following primary antibodies were incubated overnight at 4 °C in a humidified chamber: mouse anti-glial fibrillary acidic protein (GFAP) (1:500; Sigma-Aldrich), rabbit anti-laminin 1α (1:200; Abcam), rabbit anti-fibronectin (1:200; Abcam) and mouse anti-chondroitin sulfate proteoglycans (CSPGs) (1:200; Sigma-Aldrich) to evaluate scarring; rat anti-cluster of differentiation 4 (CD4) (1:500; BD biosciences), and rabbit anti-ionized calcium binding adaptor molecule 1 (Iba-1) (1:350; Wako) to evaluate the inflammatory infiltrate and rabbit anti-myelin basic protein (MBP) (1:100; Millipore) to examine demyelination. Following repeated washing steps with PBS, spinal cord sections were incubated with Alexa-labeled secondary antibodies for 1 hour at RT, namely goat anti-mouse IgG Alexa 555, goat anti-rabbit Alexa 488, goat anti-rat Alexa 488 and goat anti-mouse IgM Alexa 555 (dilution 1:250; all secondary antibodies were obtained from Invitrogen). Specificity of the secondary antibody was verified by including a control staining in which the primary antibody was omitted (data not shown). A 4,6-diamino-2-phenylindole (DAPI, Invitrogen) counterstain was performed to reveal cellular nuclei whereafter sections were mounted. Images were taken with a Nikon Eclipse 80i microscope and a Nikon digital sight camera DS-2MBWc.

Quantitative image analyses were performed on original unmodified photos using Fiji/ImageJ open source software (NIH). For standardization, analyses were performed on 5-8 spinal cord sections (per mouse) representing the lesion area, i.e. the lesion epicenter as well as consecutive sagittal sections, as previously described (12, 29, 39). T helper cell infiltration was evaluated by performing double immunostainings against CD4 and Iba-1, in order to exclude CD4+ microglial cells. All T helper cells were quantified in the entire perilesional area, extending 5 mm distal and proximal from the lesion center. These data were expressed as the average number of T cells per section. Lesion size was evaluated by delineating the area in which there was no GFAP staining. Quantification of astrogliosis (GFAP expression) and microglia/macrophage infiltration (Iba-1 expression) was performed in the perilesional area via intensity analysis within rectangular areas of 100 µm × 100 µm, extending from 600 µm cranial to 600 µm caudal from the lesion border. Demyelination was quantified by delineating the area that was absent of MBP immunoreactivity. Both the area and immunoreactivity were determined for fibronectin and laminin expression at the lesion site by delineating the area that was positive for these markers. To evaluate the expression of CSPGs, intensity of CSPG-immunoreactivity was measured perilesionally in a well-defined area surrounding the lesion center (∼200 µm thick lesion border). To maximise image readability of the representative images, the contrast and brightness of the stainings was enhanced equally in the corresponding groups (treated vs. untreated group). Furthermore, the tissue section borders were highlighted using a white line.

### Cytometric bead array – cytokine/chemokine protein levels after SCI

Cytokine/chemokine protein levels were determined locally in the injured spinal cord in mice that received either recombinant mMCP6 or a vehicle control. At 7 dpi, mice in both groups received an overdose of Nembutal and they were transcardially perfused with Ringer solution as described above. Spinal cord tissue was collected in a precisely standardized region (5 mm cranial to 5 mm caudal from the lesion center). These tissue samples were homogenized with a disposable pestle (VWR) by adding radioimmunoprecipitation assay (RIPA) lysis buffer (50 mM Tris, 150 mM NaCl, 0.5% sodium deoxycholate, 0.1% sodium dodecyl sulfate and 1.0% Triton X-100 in MilliQ), containing a complete protease inhibitor cocktail (Roche Diagnostics). Cell lysates were centrifuged at 10000 rotations per minute for 10 min and homogenates were stored at −80 °C until measurement. Total protein levels were determined using the BCA Protein Assay Kit (Thermo Fisher Scientific), according to the manufacturer’s instructions. Protein expression levels of interleukin-2 (IL-2), IL-1β, IL-13, IL-4, IL-6, IL-10, interferon gamma (IFN-γ), tumor necrosis factor alpha (TNF-α) and monocyte chemoattractant protein-1 (MCP-1) were quantitatively determined in spinal cord homogenates by flow cytometry analysis using the mouse Cytometric Bead Array (CBA) Th1/Th2/Th17 kit and specific mouse Flex set systems (BD biosciences), according to the manufacturer’s instructions. These inflammatory mediators were chosen because they are known to affect axon regeneration and other repair processes after CNS injury (38, 40, 41). Briefly, serial dilutions of the standard were prepared and samples were incubated with capture beads at RT for 1 hour. Afterwards, detection beads were added and samples were incubated for another hour at RT. Finally, all samples were washed with wash buffer and analysis was performed using a FACS Array Bioanalyzer (BD biosciences) and the FCAP software (Soft Flow Inc.).

### Culture primary OPCs and astrocytes

Primary OPCs and astrocytes were prepared from postnatal day 0 C57BL/6J mouse pups. Cortices were isolated, meninges were removed, and cells were dissociated by incubation in papain-DNAse solution (3U/ml; Sigma-Aldrich) at 37°C for 20 minutes. The mixed glial cell suspension was seeded out in poly-L-lysine (5 µg/ml) coated T75 culture flasks, containing DMEM high glucose medium (Sigma-Aldrich), supplemented with 100 U/ml penicillin and 100 µg/ml streptomycin (Invitrogen) and 10% heat-inactivated fetal calf serum (FCS) (Biowest). Cells were kept at 37°C in a humidified atmosphere of 8.5% CO_2_. OPCs were obtained on day 14 after an overnight shake-off on 250 rpm and 37°C, and were seeded (0.75 × 10^5^ cells/cm^2^) in SATO medium. Two days before treatment, OPCs were stimulated with 10 ng/ml platelet-derived growth factor-AA (PDGF-AA, Peprotech) and 10 ng/ml human fibroblast growth factor-2 (FGF-2, Peprotech) to synchronize the cells to the early OPC stage. OPCs were then treated with 500 ng/ml mMCP6 for six days, with a treatment boost every 48 hours.

Primary astrocytes were obtained from the residual attached layer after the OPC shake-off and were cultured for seven additional days. On day 21, astrocytes were seeded (0.25 × 10^5^ cells/cm^2^) in DMEM medium (+10 % FCS and 1% P/S) and stimulated with either 10 ng/ml TNF-α and 10 ng/ml IL-1β (both from Peprotech) or vehicle control. Two days later, inflammatory cytokines were washed away, and cells were treated with 500 ng/ml mMCP6 for 24 hours.

### Immunocytochemistry: OPCs and astrocytes

Primary OPCs and astrocytes were fixed with 4% PFA for 30 min at RT. The cells were blocked and permeabilised with 0.1% Triton X-100 in 1% bovine serum albumin (BSA) for 30 min, followed by 4 hour incubation with mouse anti-O4 IgM (1:1000; R&D Systems) and rat anti-MBP (1:500; Serotec) antibodies (OPCs) or 2 h incubation with mouse anti-GFAP (1:200, Sigma) (astrocytes). Next, cells were washed in PBS and incubated with Alexa fluor secondary antibodies (Invitrogen) for 1 hour. Nuclei were counterstained with 1 µg/ml DAPI. Coverslips were mounted with DAKO mounting medium (Dako) and analyzed using a Leica DM2000 LED fluorescence microscope. Image quantification was carried out using Fiji, ImageJ software.

### Statistical analysis

All statistical analyses were performed using GraphPad Prism 5.01 software (GraphPad Software, Inc.). Data sets were analyzed for normal distribution using the D’Agostino-Pearson normality test. Functional recovery *in vivo*, histological analyses of GFAP and Iba-1 immunoreactivity were statistically analyzed using two-way analysis of variance (ANOVA) for repeated measurements with a Bonferroni *post hoc* test for multiple comparisons. All other differences between two groups were evaluated using the nonparametric Mann-Whitney U test. Data were presented as mean + standard error of the mean (SEM). Differences were considered statistically significant when p < 0.05.

## Results

### Recombinant mMCP6 improves hind limb motor recovery after traumatic SCI

Previously, we showed that MCs have beneficial effects on the functional outcome after SCI partly via the scar remodellig properties of endogenous mMCP6 (30). As a follow-up study, we here aimed to determine whether recombinant mMCP6 can be applied therapeutically to improve recovery after traumatic SCI. For this purpose we developed mMCP6 in-house, and first determined the proteolytic stability of the protease under physiological conditions. We found that the tryptase activity of mMCP6 rapidly decreased at pH 7.4, with a half-life of approximately 1.5 hours (**Fig. 1B**). Therefore, we decided to adminster recombinant mMCP6 daily in the subacute phase after injury via an intrathecal catheter system to ensure ‘constant’ proteolytic levels at the lesion site (**Fig. 1A**). An illustration of the experimental approach is presented in **Fig. 1C**. WT C57BL/6j mice were subjected to a dorsal T-cut hemisection injury and they were assigned to a treatment group, receiving either recombinant mMCP6 or a proteolytic inactive vehicle, which was prepared by heating recombinant mMCP6 to 100°C. Both groups experienced severe hind limb paralysis in the acute phase (0 dpi – 2 dpi) after injury, and mice in both treatment groups gradually improved over time (**Fig. 1D**). However, from day 7 onwards until the end of the observation period (27 dpi), locomotor function was significantly improved in the mMCP6-treated group compared to the control group (**Fig. 1D**). Individual BMS scores at 27 dpi are shown for both treatment groups in **Fig. 1E**.

### Local delivery of recombinant mMCP6 did not reduce scar formation at the lesion center after SCI

Using knockout mice, we previously showed that the absence of endogenous mMCP6 results in excessive scarring at the lesion site after SCI (30). This scarring response is characterized by the deposition of axon-growth inhibitory CSPGs in the perilesional area (8, 42), and the formation of a fibrous and basement membrane-rich compartment at the lesion center (13, 43, 44). Here, we investigated whether the therapeutic effect of recombinant mMCP6 was also mediated via scar remodeling. We did not observe an effect of mMCP6 on the fibrous scarring response. Both the area (**Fig. 2Ai/ii, B**) and immunoreactivity (**Fig. 2Ai/ii, C**) of fibronectin were similar between the mMCP6-treated and the control group at the lesion center. In addition, the area and immunoreactivity of the basement membrane component laminin (**Fig. 2Aiii/iv, D, E**), was comparable between the two treatment groups after injury. The intensity of axon-growth inhibitory CSPGs was analyzed in a well-defined area around the lesion center (white encircled area in **Fig. 2Av/vi**). Immunoreactivity for CSPGs was also not significantly different between the control group and the mMCP6-treated group in the perilesional area (**Fig. 2Av/vi, F**).

**Figure 2.**
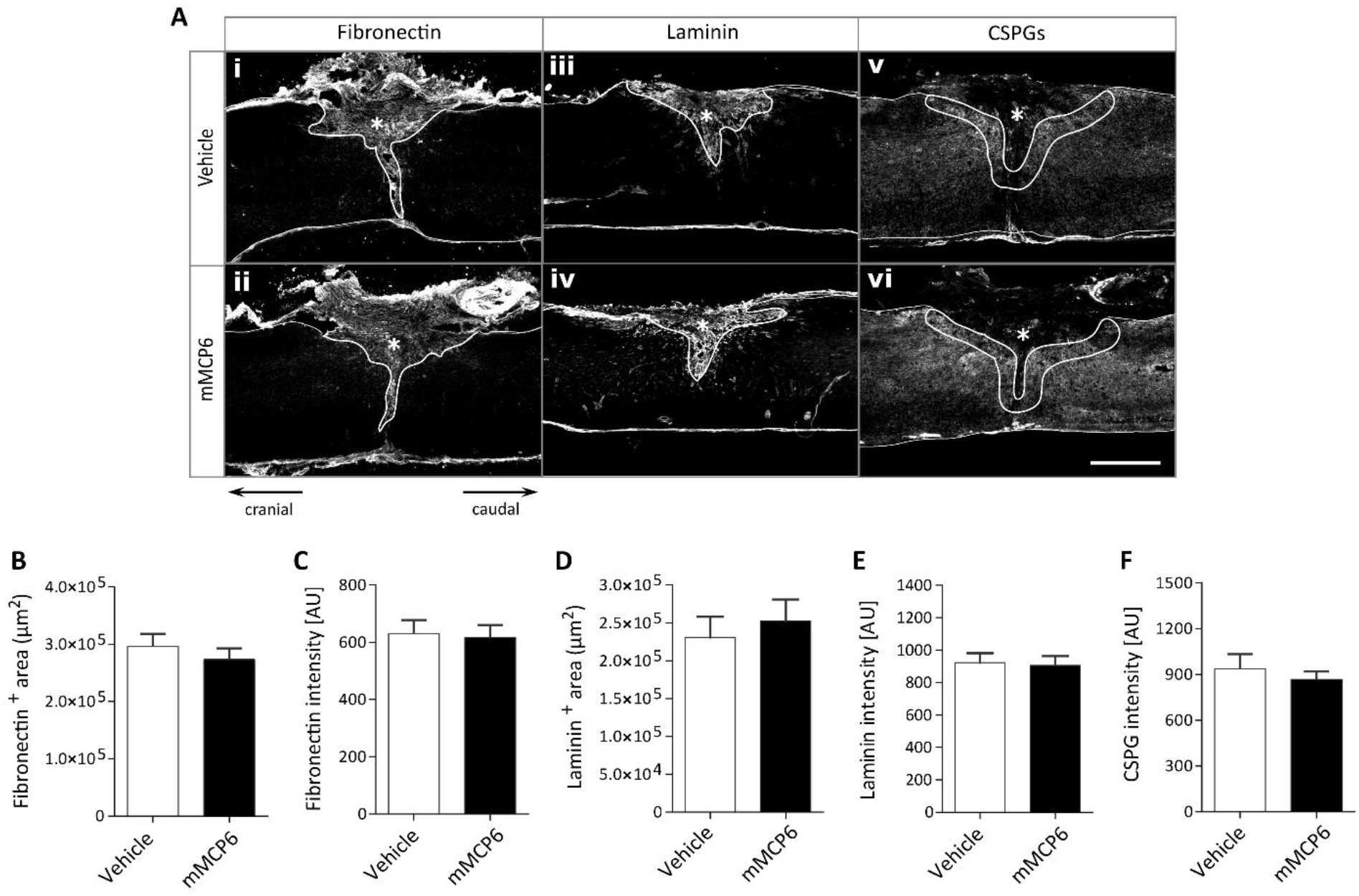
Local administration of recombinant mMCP6 does not affect the scarring response after SCI. **(A)** Representative photomicrographs of fibronectin, laminin and CSPG expression at the lesion center for the vehicle control group and mMCP6-treated group, respectively. **(B, C)** Immunofluorescence for fibronectin was chosen to analyze the scarring response after injury (area marked by white line in Ai/ii). Image analysis revealed that both the (B) area and (C) immunoreactivity for fibronectin at the lesion center were not significantly different between the (Ai) control group and the (Aii) mMCP6-treated group. **(D, E)** Both the (D) area and (C) immunoreactivity for laminin were also not significantly different between the (Aiii) control group and the (Aiv) mMCP6-treated group at the lesion site. **(F)** Immunoreactivity for CSPGs was determined in the perilesional area (area marked by white encircled line in Av/vi). No difference in CSPG intensity was observed between the (Av) vehicle group and the (Avi) mMCP6-treated group, respectively. Data are presented as mean values + SEM. mMCP6: n = 12; vehicle: n = 11. Asterisks indicate the lesion center. Scale bar = 500 µm.

### Locally applied recombinant mMCP6 did not change the inflammatory response after SCI

Injury to the spinal cord is always associated with an inflammatory response, which can have both detrimental and beneficial effects on axon regeneration and recovery processes (21, 38). This inflammatory response is characterized by the activation and infiltration of immune cells, and their secretion of inflammatory mediators at the lesion site (11, 45). In a next step, we determined whether local application of recombinant mMCP6 influenced (1) the cellular immune response and/or (2) the cytokine/chemokine expression levels at the lesion site after SCI. Immunohistochemical assessment of Iba-1 and CD4 markers were chosen to investigate the infiltration of microglia/macrophages and T helper cells, respectively, into the lesion area. This revealed that Iba-1 immunoreactivity peaked in close proximity to the lesion area and declined at further distances from the lesion site in both groups. However, no significant difference was observed between the control group (**Fig. 3Ai, B**) and the mMCP6-treated group (**Fig. 3Aii, B**). In addition, the infiltration of CD4^+^ T cells into the lesion area was not significantly different between the two groups (**Fig. 3Aiii/iv, C**). A CBA analysis was performed to examine cytokine/chemokine protein levels in the injured spinal cord at 7 dpi. This time point was chosen because the difference in functional recovery between the mMCP6-treated and vehicle control group starts at this point (**Fig. 1D**). IL-1β, TNF-α and MCP-1 protein levels were upregulated after injury, but there was no difference between the mMCP6-treated group and the vehicle control group (**Fig. 3D**). In contrast, local IL-2, IL-4, IL-6, IL-10, IL-17α and IFNγ protein levels were not elevated after injury and furthermore, no significant difference was observed between the two treatment groups at 7 dpi (**Fig. 3D**).

**Figure 3.**
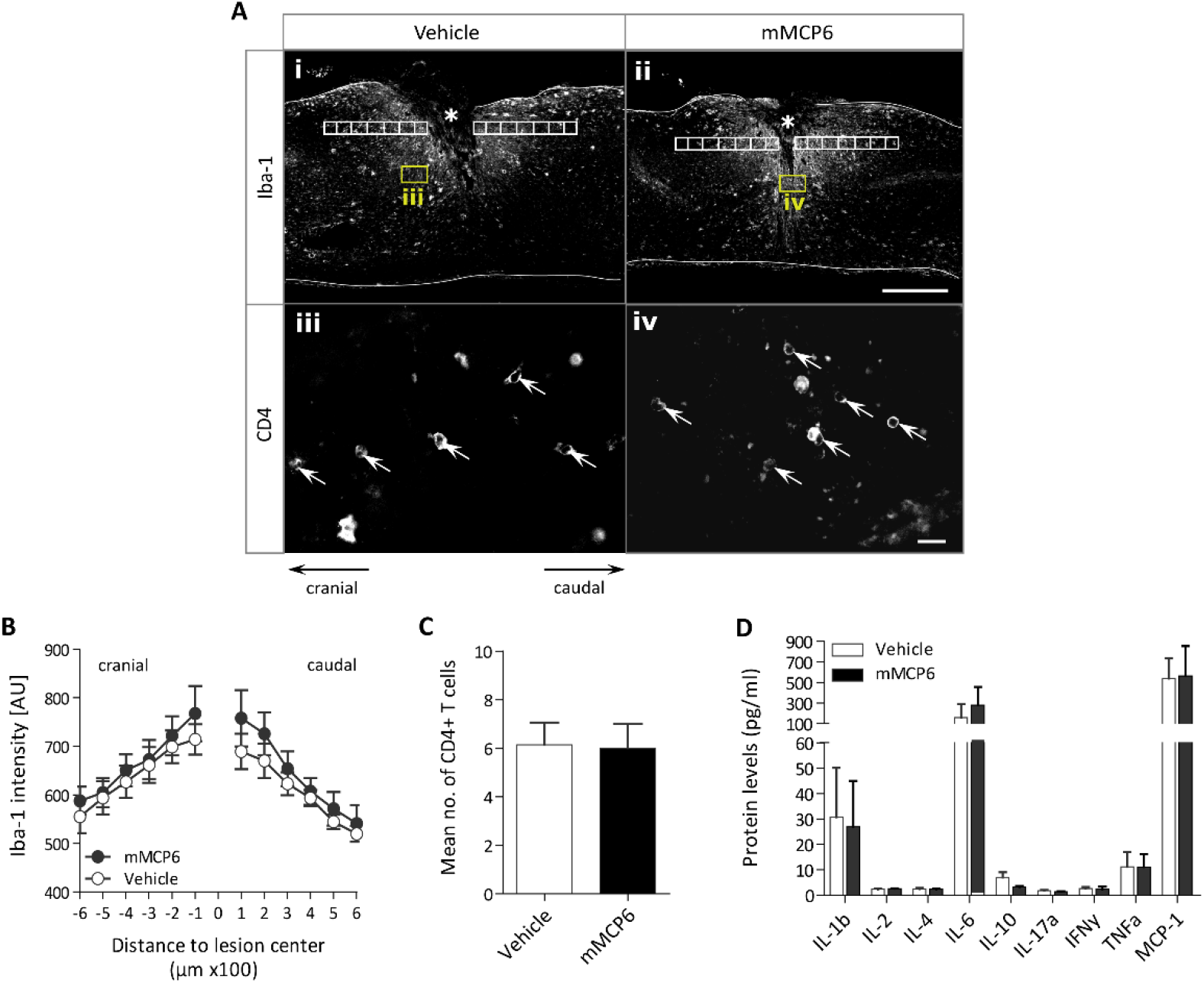
Administration of recombinant mMCP6 did not change inflammatory processes after SCI. **(A)** Representative photomicrographs of Iba-1 and CD4 expression at the lesion site for the vehicle control group and mMCP6-treated group, respectively. Boxed regions in Ai and Aii (Iba-1 staining) are shown in Aiii and Aiv (CD4 staining) to localize the T cells in the spinal cord. **(B)** Iba-1 immunoreactivity was quantified within square areas of 100 μm × 100 μm (white boxes in Ai/ii), extending from 600 μm cranial to 600 μm caudal from the lesion border. No difference in Iba-1 immunoreactivity was observed in the perilesional area between the (Ai) control group and the (Aii) mMCP6-treated group. **(C)** In addition, immunofluorescence for CD4 showed that the number of infiltrating T helper cells into the lesion area was not significantly different between the (Aiii) vehicle group and the (Aiv) mMCP6-treated group. White arrows in Aiii/iv indicate CD4+ T cells. Scale bars: Ai/ii = 500 μm and Aiii/iv = 20 μm. Data were presented as mean ± SEM. n = 10-12/group. **(D)** Cytometric Bead Array (CBA) analysis were performed to examine cytokine/chemokine protein levels at the lesion site between the control group and the mMCP6-treated group at 7 dpi. No difference in protein levels of IL-1β, IL-2, IL-4, IL-6, IL-10, IL-17α, IFNγ, TNFα and MCP-1 was observed between the two treatment groups at 7 dpi. Data are presented as mean values + SEM. n = 6/group.

### Recombinant mMCP6 does not affect serotonergic fiber growth after SCI

To test for beneficial effects of recombinant mMCP6 on axon regeneration after a spinal cord hemisection injury, we visualized serotonergic raphespinal projections by an anti-serotonin (5HT) antibody. We determined 5HT^+^ fiber growth (number and total length) caudal from the lesion site in the vehicle control group and the mMCP6-treated group after SCI (**Fig. 4**). Representative immunofluorescence images and Neurolucida-like drawings of 5HT^+^ fiber growth are illustrated for the control group (**Fig. 4Ai/iii**) and mMCP6-treated group (**Fig. 4Aii/iv**), respectively. We observed a slight, non-significant, trend towards an increase in the number and total length of 5HT^+^ fibers in the mMCP6-treated group compared to the vehicle control group (**Fig. 4Ai-iv, B, C**).

**Figure 4.**
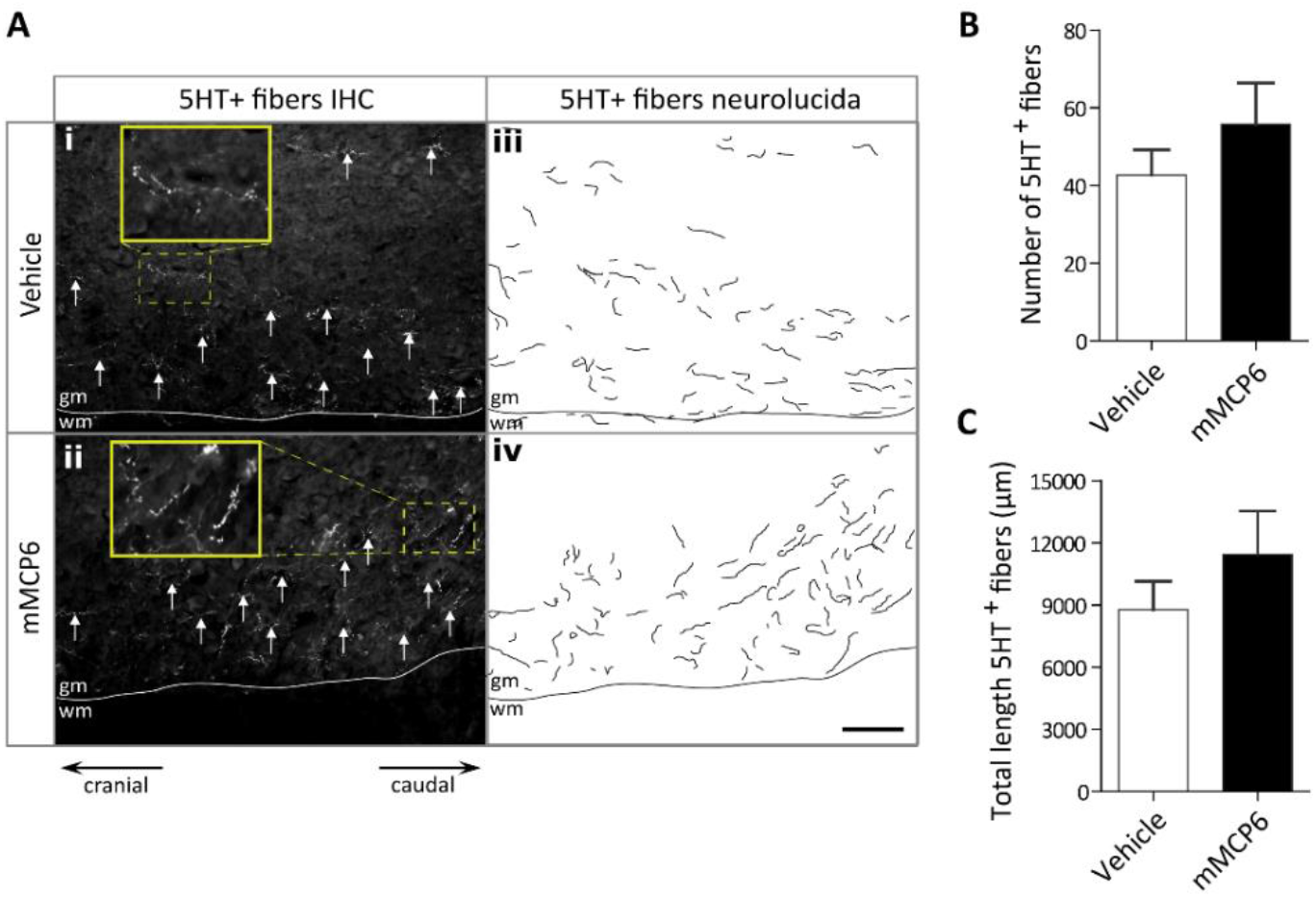
Recombinant mMCP6 does not significantly improve raphespinal axon regeneration after SCI. **(A)** Representative sagittal sections of the 5HT immunofluorescence staining are shown for (Ai) control mice and the (Aii) mMCP6-treated mice, respectively. White arrows indicate 5HT+ fibers. Neurolucida-like drawings of 5HT-immunoreactive fibers are also illustrated for the (Aiii) vehicle control group and the (Aiv) mMCP6-treated group. **(B, C)** Fiber analysis with ImageJ showed a slight trend towards an increase in the (B) number and (C) total length of 5HT+ fibers caudal from the lesion site in the mMCP6-treated group. Data are presented as mean values + SEM. Dashed lines in (A) mark the border between the grey matter (gm) and white matter (wm). Scale bar = 500 µm. n = 12-14 sections/group.

### Local administration of mMCP6 reduces astrogliosis and the demyelinated area after SCI

To further characterize the therapeutic effect of recombinant mMCP6, we investigated whether the improvement in functional recovery was associated with a less severe lesion pathology. Immunofluorescence labelling for glial fibrillary acidic protein (GFAP) revealed no difference in lesion size between the mMCP6-treated group and the vehicle control group (**Fig. 5Ai/ii, B**). Astrogliosis was determined by measuring GFAP intensity in the perilesional area (white boxes in **Fig. 5Aiii/iv**). GFAP intensity peaked in both treatment groups at the lesion border and decreased further caudally and cranially from the lesion site (**Fig. 5Aiii/iv, C**). Astrogliosis was significantly decreased in the mMCP6-treated mice compared to the control mice at 27 dpi (**Fig. 5Aiii/iv, C**). Immunofluorescence labelling for myelin basic protein (MBP) revealed that the demyelinated area was significantly reduced in the mMCP6-treated group compared to control mice (**Fig. 5Av/vi, D**).

**Figure 5.**
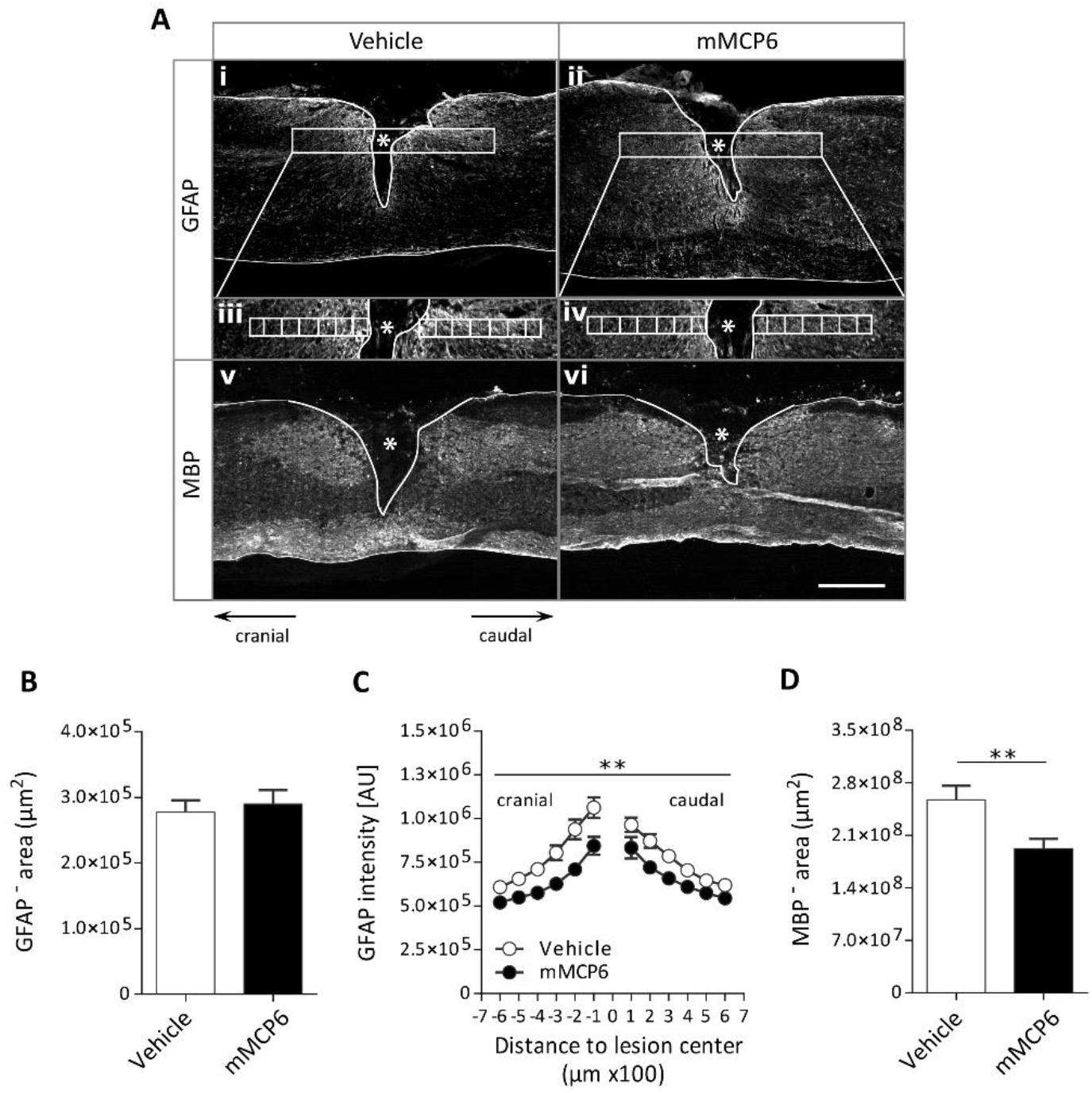
Local application of recombinant mMCP6 reduces astrogliosis and the demyelinated area after SCI. **(A)** Representative photomicrographs of the (Ai/ii) lesion size, (Aiii/iv) astrogliosis and (Av/vi) demyelinated area for control and mMCP6-treated mice after SCI. **(B)** Immunofluorescence for glial fibrillary acidic protein (GFAP) was chosen to determine the lesion size (GFAP-negative area, marked by white line in Ai/ii). No difference in lesion size was observed between the (Ai) control group and the (Aii) mMCP6-treated group after injury. **(C)** Astrogliosis was quantified by measuring GFAP intensity within square areas of 100 μm × 100 μm (white boxes in Aiii/iv), extending from 600 μm cranial to 600 μm caudal from the lesion border. Astrogliosis was significantly decreased in the (Aiv) mMCP6-treated group compared to the (Aiii) control group. Images Aiii/iv are magnifications of the white rectangular boxes in Ai/ii, respectively. **(D)** Immunofluorescence for myelin basic protein (MBP) was used to characterize the demyelinated area (MBP-negative area, marked by white line in Av/vi). A significant reduction in demyelination (or alternatively improvement in remyelination) was observed in the (Avi) mMCP6-treated group compared to the (Av) control. Data are presented as mean + SEM. vehicle: n = 11; mMCP6: n = 12. Scale bar in Ai, ii, v, vi = 500 µm. ** p < 0.01

### Recombinant mMCP6 boosts OPC differentiation and reduces astrogliosis *in vitro*

As a final step in the study, we aimed to investigate how recombinant mMCP6 could mechanistically reduce astrogliosis and improve myelination levels *in vivo* after SCI. To this end, we set up *in vitro* experiments; firstly primary oligodendrocyte precursor cells (OPC) cultures were stimulated with recombinant mMCP6. We found that mMCP6 stimulated the differentiation towards a more mature phenotype, as indicated by increased O4 (premature oligodendrocyte marker) and MBP (mature myelinating glia) expression levels in the mMCP6-treated cell cultures compared to the vehicle control (**Fig. 6Ai-iv, B, C**). Secondly, astrocytes, either controls or activated via the inflammatory cytokines TNF-α and IL-1β (mimicking the inflammatory conditions seen in SCI), were treated with recombinant mMCP6 in culture. We observed that recombinant mMCP6 significantly reduced astrogliosis as indicated by lower GFAP expression levels per cell, in both the control condition (**Fig. 6Di/ii, E**) and also inflammation-induced astrogliosis (**Fig. 6Diii/iv, F**).

**Figure 6.**
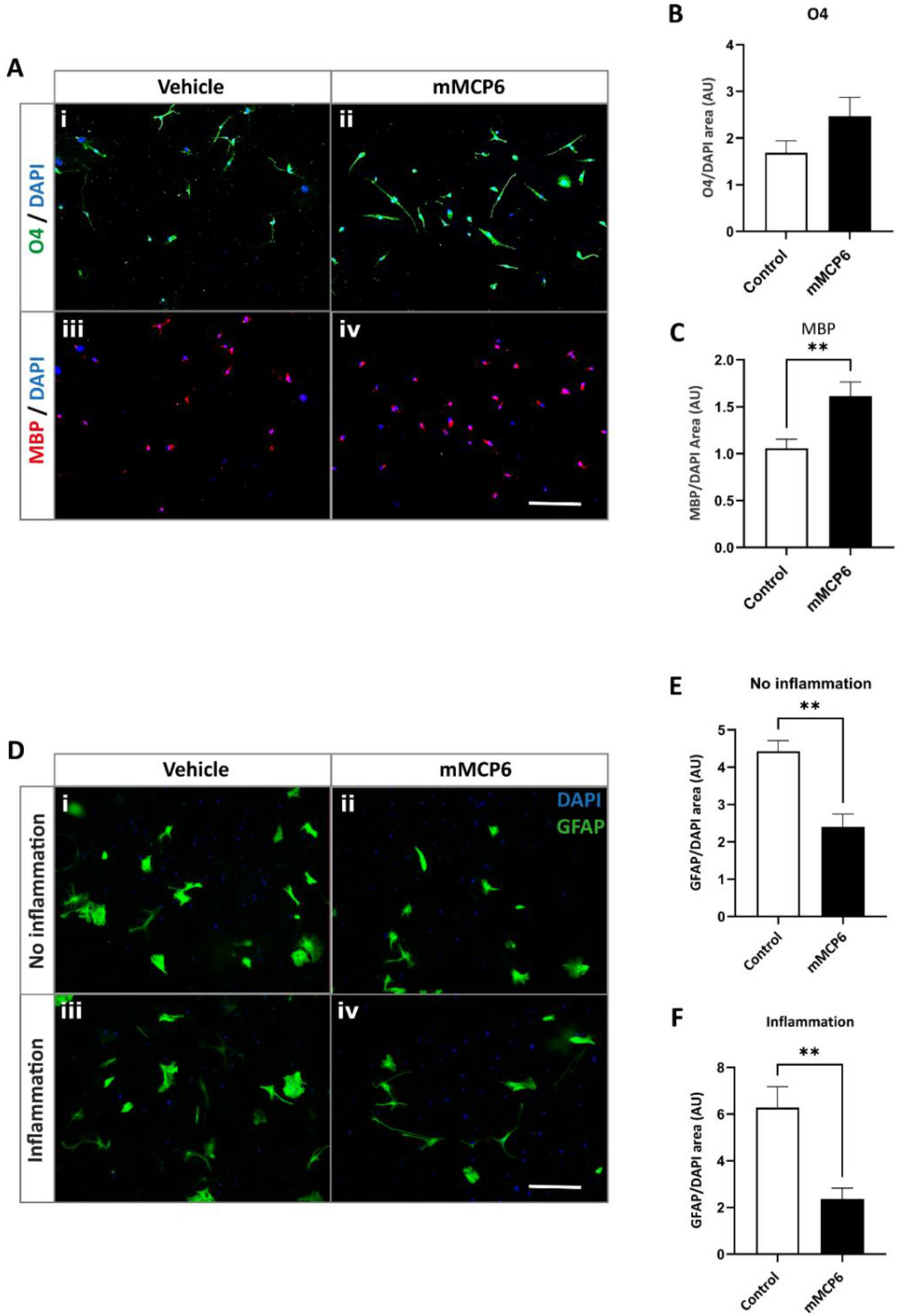
Recombinant mMCP6 significantly stimulates OPC differentiation towards MBP-expressing cells and reduces astrogliosis *in vitro*. (A-C) Primary OPCs were treated with 500 ng/ml mMCP6 or its inactive form as a control. Cells were kept in culture for six days (treatment boost was performed at day 0, day 2 and day 4). **(A)** The effect on cellular differentiation was assessed via immunofluorescence against O4 (premature oligodendrocyte marker, in green, Ai/ii) and MBP (differentiated oligodendrocytes, in red, Aiii/iv). **(B, C)** Integrated density analysis indicate increased expression levels of O4 (B) and MBP (C) in mMCP6-treated OPC cultures, compared to controls. (D-F) Primary astrocytes, either unstimulated or primed with 10 ng/ml TNF-α/IL-1β (inflammation-triggered astrogliosis), were also stimulated with recombinant mMCP6 (500 ng/ml) or an inactive vehicle for 24 hours. **(D)** The effect on astrocyte activation was examined via immunofluorescence against GFAP (in green, Ai-iv), and **(E, F)** image analysis showed that mMCP6 significantly reduced astrogliosis in culture. Nuclei (in blue in A and D) were visualized using DAPI. Scale bar in A and D: 50 µm. Data are represented as mean values + SEM. Mann-Whitney U-test.

## Discussion

To date, no effective clinical treatment exists that can restore lost functions after traumatic SCI. Several pathophysiological processes (e.g. inflammation, scarring, demyelination) contribute to the impaired functional outcome after SCI and it is therefore likely that a combination of therapies will be necessary to improve recovery (1, 4, 20). Previous research indicates that MCs protect the CNS after traumatic injury via several mechanisms, namely by suppressing harmful inflammatory reactions via mMCP4 that cleaves pro-inflammatory mediators (28, 29) and by reducing scar tissue formation, mainly via mMCP6 (30). In addition, mMCP4 has anti-scar properties (46). These findings indicate that endogenous mMCP4 and mMCP6 support repair processes after CNS injury. It is therefore, tempting to speculate that a combination of both factors will have a synergistic therapeutic effect on functional outcome after CNS trauma. In the present study, we addressed whether mMCP6 can be used therapeutically to improve recovery after SCI. Several other studies have shown that extracellular proteases, such as MMPs and plasminogen activators, can exert both detrimental and beneficial effects on regenerative processes in the CNS, depending on the time point after injury (47, 48). In the acute phase, they disrupt the blood brain barrier (BBB), thereby supporting pro-inflammatory processes that induce secondary tissue damage (49-51). However, at later time points, extracellular proteases can improve recovery by stimulating axon growth, synaptic plasticity, remyelination, and angiogenesis (52-55). Hence it is essential to define the ‘optimal’ therapeutic time window, and we decided to apply mMCP6 daily from 4 dpi until 10 dpi (i.e., the subacute phase) via an intrathecal catheter. Drug delivery in the intrathecal space is an effective way to distribute drugs, locally and in a time-dependent manner to the injured CNS (56-58). By following this therapeutic approach, we found that the functional outcome was significantly improved in mice that received recombinant mMCP6 compared to the vehicle-treated group. Moreover, both the demyelinated area and astrogliosis in the perilesional area were significantly reduced in mMCP6-treated mice. In addition, we show that serotonergic fiber growth behind the lesion site tends to improve in mice that received recombinant mMCP6. Surprisingly, we did not observe any effect of mMCP6 on the lesion size or the scarring response after SCI, which is in contrast to our previous findings in mMCP6 knockout mice (30). Moreover, expression levels of the axon-growth inhibitory CSPGs and the ECM components fibronectin and laminin were comparable between both groups at 28 dpi. These data suggest that additional and more complex mechanisms are involved via which recombinant mMCP6 improves recovery after SCI.

Astrocytes are a major type of glial cell in the CNS, and they play an essential role in pathophysiological conditions such as trauma. In response to SCI, astrocytes undergo characteristic phenotypic changes known as reactive astrogliosis, which can exert both beneficial and detrimental effects on the surrounding lesion microenvironment (8, 59-61). The activation state of astrocytes (different phenotypes), the time point after injury, as well as the distance of these astrocytic populations from the lesion site all determine their permissive or inhibitory behavior after SCI (62). Reactive astrocytes located in the perilesional area are considered to be detrimental in the subacute to chronic phase after SCI, where they create a physical barrier for regenerating axons and their release of chemical mediators such as CSPGs, TNF-α, and MMP9, block repair processes or cause secondary tissue damage (8, 63). Some studies show that tryptase activates astrocytes via a protease activated receptor 2 (PAR-2) dependent mechanism (64), but it is unclear how mMCP6 can suppress astrogliosis as observed in our study. Injury to the spinal cord mobilizes astrocyte-reactivating factors, including ATP, endothelin 1 (ET1), glutamate and several inflammatory cytokines such as IL-1β, TNF-α, IL-6, LIF and oncostatin M, but mechanical stress can also induce astrocyte reactivity (65, 66). Given the broad-substrate cleavage capacity (32), it is possible that recombinant mMCP6 indirectly suppresses perilesional astrogliosis by cleaving and inhibiting these factors. Indeed, we show decreased astrogliosis after recombinant mMCP6 treatment *in vitro*, where astrocytes have been incubated with TNF-α/IL1-β to induce an inflammatory phenotype.

Astrocytes also play an important role in regulating de- and remyelination processes. Some studies have shown that reactive astrocytes impair remyelination in several demyelinating pathologies such as multiple sclerosis and SCI (67-70). Oligodendrocyte loss and myelin degradation take place primarily during the subacute and chronic phase of SCI (17, 22, 71, 72). Demyelination results in less efficient signal transduction, and chronically demyelinated axons are more vulnerable to degenerative processes (73). Astrocytes secrete several factors that can induce demyelination or inhibit the differentiation of oligodendrocyte progenitor cells (OPCs) into myelin-forming oligodendrocytes, such as ET-1 (74). Moreover, it has been shown that remyelination is more efficient after transplantation of neonatal OPCs into an astrocyte-free area of demyelination (75). This may explain why the reduction in astrogliosis in mMCP6-treated animals is accompanied by a decrease in demyelinated area. Alternatively, it is possible that mMCP6 has a positive effect on the remyelination process after SCI by facilitating the differentiation of the OPCs towards mature myelinating oligodendrocytes. Therefore, we have assessed the effects of recombinant mMCP-6 on OPC differentiation *in vitro*, which indeed indicated increased numbers of differentiated MBP^+^ oligodendrocytes after incubation of OPCs with mMCP6.

In conclusion, these data show for the first time that recombinant mMCP6 can be used therapeutically to improve recovery after SCI. In contrast to our previous findings in mMCP6 knockout mice, scarring was not altered, suggesting that additional and more complex mechanisms involving reduced astrogliosis and demyelination, or improved remyelination by increased oligodendrocyte differentiation stimulate recovery after SCI.

## Ackowledgments

The authors thank Dr. Leen Timmermans (Hasselt University) for her technical assistance and help with the CBA protein analysis and immunohistochemistry. This study was supported in part by grants from Fonds voor Wetenschappelijk Onderzoek-Vlaanderen (FWO-Vlaanderen) to SH (G067715N, G091518N, G0C2120N) and from Agentschap voor Innovatie door Wetenschap en Technologie (IWT) to TV (101517).

## Author contribution

T.V. contributed to experimental design, data collection, data interpretation and manuscript writing. S.L. and N.G. contributed to data collection, data interpretation and revision of the manuscript. A.T. and T.V. contributed to data collection and revision of the manuscript. D.D. & G.P. contributed to experimental design and revision of the manuscript. S.H. contributed to experimental design, data interpretation and reviewed the manuscript. All authors discussed the results and contributed significantly to the final version of the manuscript.

## Data availability statement

All data supporting the findings of this study are available from the corresponding author on reasonable request.

